# Microbial Cells Harboring a Mitochondrial Gene Are Capable of CO2 Capture

**DOI:** 10.1101/411090

**Authors:** Yanchao Zhou, Lan Ouyang, Xiao Yi, Tao Gan, Jinhuan Qi, Yulin Wan, Yuchuan Wang, Shanshan An, Yunfan Shi, Wei Yang, Wenze Chen, Zhiyao Luo, Jing Li, Jun Luo, Xiren Nuertai, Xiang Zhu, Fan Yang, Beibei Zhao, Weiwei Zhang, Zi-Wei Ye, Xiaoxiao Zhang, Shaoping Weng, Qiuyun Liu, Weiguo Cao, Jianguo He

## Abstract

Global warming is escalating with increased temperatures reported worldwide. Given the enormous land mass on the planet, biological capture of CO_2_ remains a viable approach to mitigate the crisis as it is economical and easy to implement. In this study, a gene capable of CO_2_ capture was identified via selection in minimal media. This mitochondrial gene named as *OG1* encodes the OK/SW-CL.16 protein and shares homology with cytochrome oxidase subunit III of various species and PII uridylyl-transferase from *Loktanella vestfoldensis* SKA53. CO_2_ capture experiments indicate that δ^13^C was substantially higher in the cells harboring the gene *OG1* than the control in the nutrition-poor media. This study suggests that CO_2_ capture using engineered microorganisms in barren land can be exploited to address the soaring CO_2_ level in the atmosphere, opening up vast land resources to cope with global warming.

**IMPORTANCE:** Global warming crisis is deteriorating with increased CO2 levels in the atmosphere each year. Action must be taken before catastrophic consequences occur in the not-so-distant future. Biological capture of CO2 is a feasible approach to alleviate the current crisis. We have identified a mitochondrial gene which demonstrated CO2 utilization capability. Data presented in this study suggest that CO2 capture using engineered microorganisms can be harnessed to address the ever-rising CO2 level in the atmosphere.

---

It is widely believed that the combustion of fossil fuel contributes to the rising CO_2_ level in the atmosphere, by a surge of estimated 100 ppm since its pre-industrial level^1^. Global temperature has risen by more than half a degree in the past 50 years as a result of greenhouse gas emissions. Current technologies for carbon capture and storage and chemical fixation of CO_2_ are varied^2^. The amount of CO_2_ captured relative to the magnitude of CO_2_ emitted and energy required are two major concerns. From this point of view, biological CO_2_ capture is a viable approach with minimal energy demand. Here we describe the cloning of a mitochondrial gene which is able to capture CO2, and the engineered microbes showed great potential in addressing the global warming issues.

## Results

To search for genes that can capture CO_2_, we constructed and screened a random DNA library. The effort has ended with a sequence from an unknown DNA library, which shares homology with the genome of Homo sapiens haplogroup H63 mitochondrion (Supplementary Fig. S1) and gave rise to *Escherichia coli* colonies in nitrogen free media. More importantly, these colonies were later shown to be able to grow on carbon deficient media. Supplement in rich media Luria Broth with vitamin C was essential for the successful sequencing of the positive clone pOG, suggesting that the OG-encoded protein might catalyzes a stressful oxidative reaction. The protein encoded in the subclone pOG1 is identical to a known protein OK/SW-CL.16^3^, which shares homology with PII uridylyl-transferase from *L. vestfoldensis* SKA53 and cytochrome oxidase subunit III from *Macaca mulatta* (Supplementary Fig. S2). The OK/SW-CL.16 protein harbors a class I PxxP motif, a predicted actinin-interacting region, and a RIM1-like sequence^3^. Previous study has shown that OK/SW-CL.16 acted as a binding partner for actinin-4^3^.

Yeast extract was included in carbon-free media to allow proton traffic and attenuate osmotic pressure since its peptide and protein constituency are rich in charged amino acids and hydrogen bond donors and acceptors (Supplementary Fig. S3 and S4). Without supplement of minimal amount of proteins and peptides, pOG1 grew very poorly in carbon free nutrition-poor liquid media (Supplementary Fig. S3).

The commercially available ^13^C-Urea Breath Test was based on the release of CO_2_ by urease^4,5^. It was adapted in this study to investigate the potential capacity of CO_2_ capture using our clones. Isotopic measurements at the end of carbon fixing experiments (Fig. 1; Supplementary Tables S1-S8) indicate that δ^13^C was substantially different between the clones harboring OG1 gene and the control clone in both *E. coli* and *Saccharomyces cerevisiae* (Fig. 1A and 1D), suggesting that the clones harboring OG1 gene may have higher assimilation rate toward CO_2_. Since microbial cells were submerged in 50 ml liquid media without shaking, it suggests that engineered microbial cells might be able to capture CO_2_ under surface soil. ATP supplement diminished the CO_2_ fixing activity of the pOG1 clone (Fig. 1A). Although a clone pOGDR1 obtained from directed evolution had significantly higher δ^13^C than pOG1 clone at pH 8.0, its δ^13^C value was lower than that of pOG1 clone at pH 7.0 (Fig. 1G and 1J; Table 1), suggesting that proton traffic may be integral in CO2 capture. Treatments with the carbonic anhydrase inhibitor acetazolamide generated no difference in δ^13^C patterns between pOG1 groups and pSK- groups (p=0.567; Fig. 2, S9 and S10), suggesting that spontaneous hydration of CO2 was present. Alternatively, the *E. coli* carbonic anhydrase may be insensitive to acetazolamide. *OG1* was highly expressed in engineered *E. coli* host cells, which was determined via Realtime fluorescent quantitative PCR (Fig. 3; Supplementary Tables S11 and S12).

**Figure 1.**
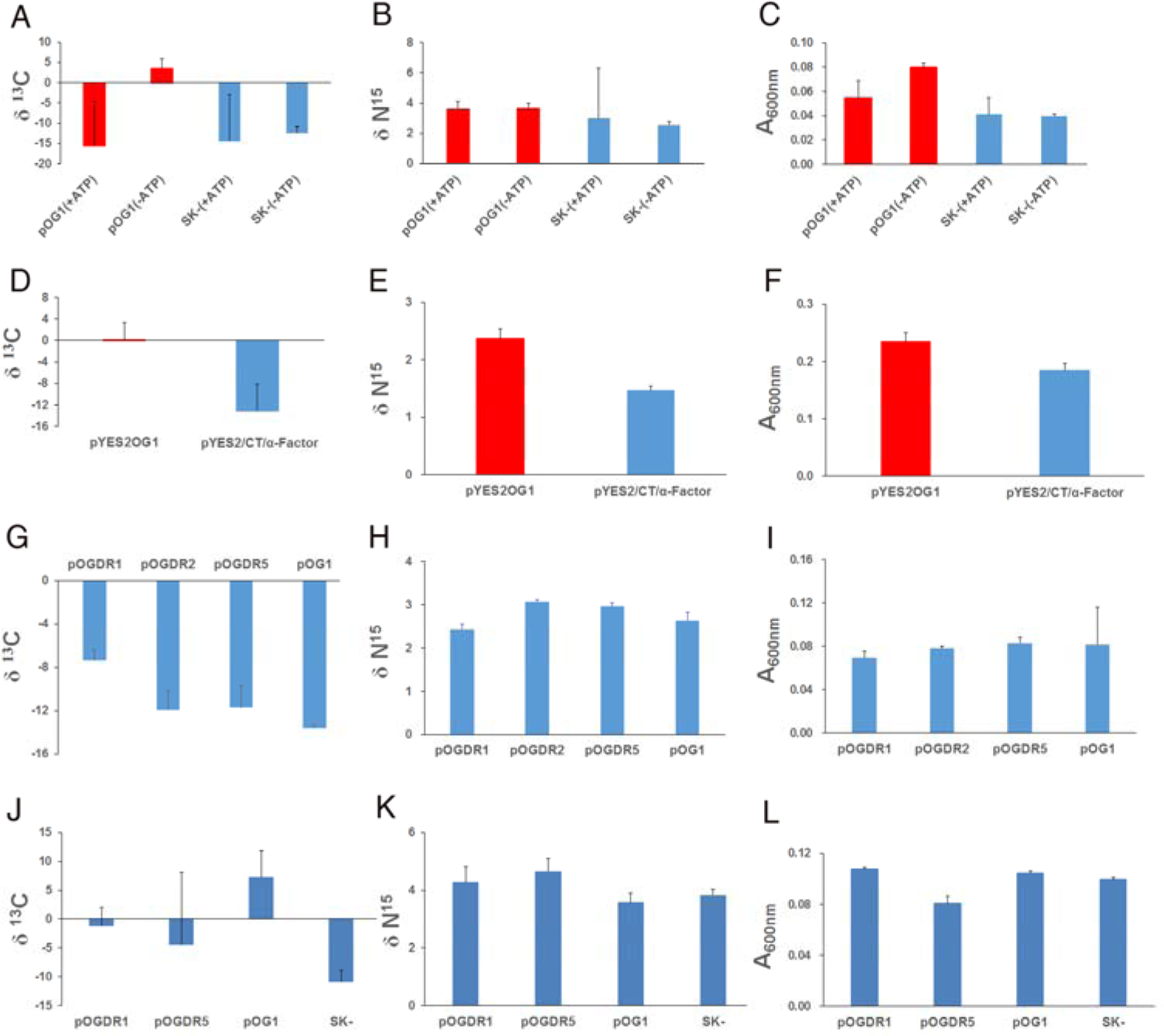
δ^13^C and δN^15^ for CO_2_ capture experiments. **A,B,C.** Effects of ATP usage on δ_13_C, δN^15^, OD600nm post-carbon capture. Orders of the rest of the panels are likewise. **D.** δ_13_C with *S. cerevisiae* clones. **G.** δ_13_C by clones obtained via directed evolution. From A to I, all were propagated in media at pH 8.0. **J.** δ_13_C from clones propagated in media at pH 7.0. Statistical evaluations for ^13^C are as follows (univariate General Linear Model, two tailed, and n = 3 unless specified): **A.** 0.041 (pOG1: ATP vs absence of ATP). **D.** 0.001 (pYES2OG1 vs pYES2/CT/α-Factor; n=5). **G.** 0.012 (pOG1 vs pOGDR1; Games-Howell post hoc tests). **J.** 0.032 (pOG1 vs pSK-; Games-Howell post hoc tests), 0.047 (pSK- vs pOGDR1; Games-Howell post hoc tests). The data were presented as average values with one standard deviation.

**Figure 2.**
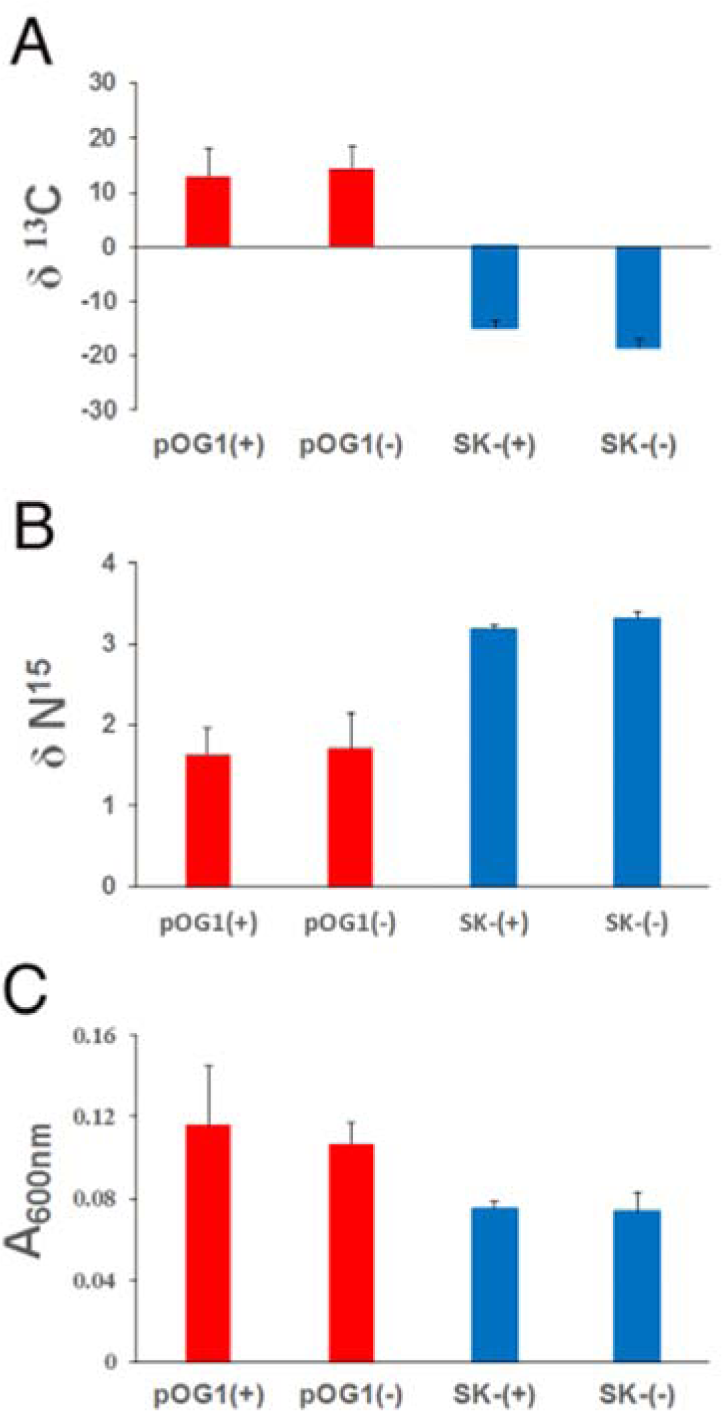
A, B, C. Effects of acetazolamide usage on ^13^C, δN^15^, OD600nm post-carbon capture. Treatments with carbonic anhydrase inhibitor acetazolamide (n=3). (+): Presence of 50 µg/ml acetazolamide in the media. (−): No acetazolamide in the media. The data were presented as average values with one standard deviation.

**Figure 3.**
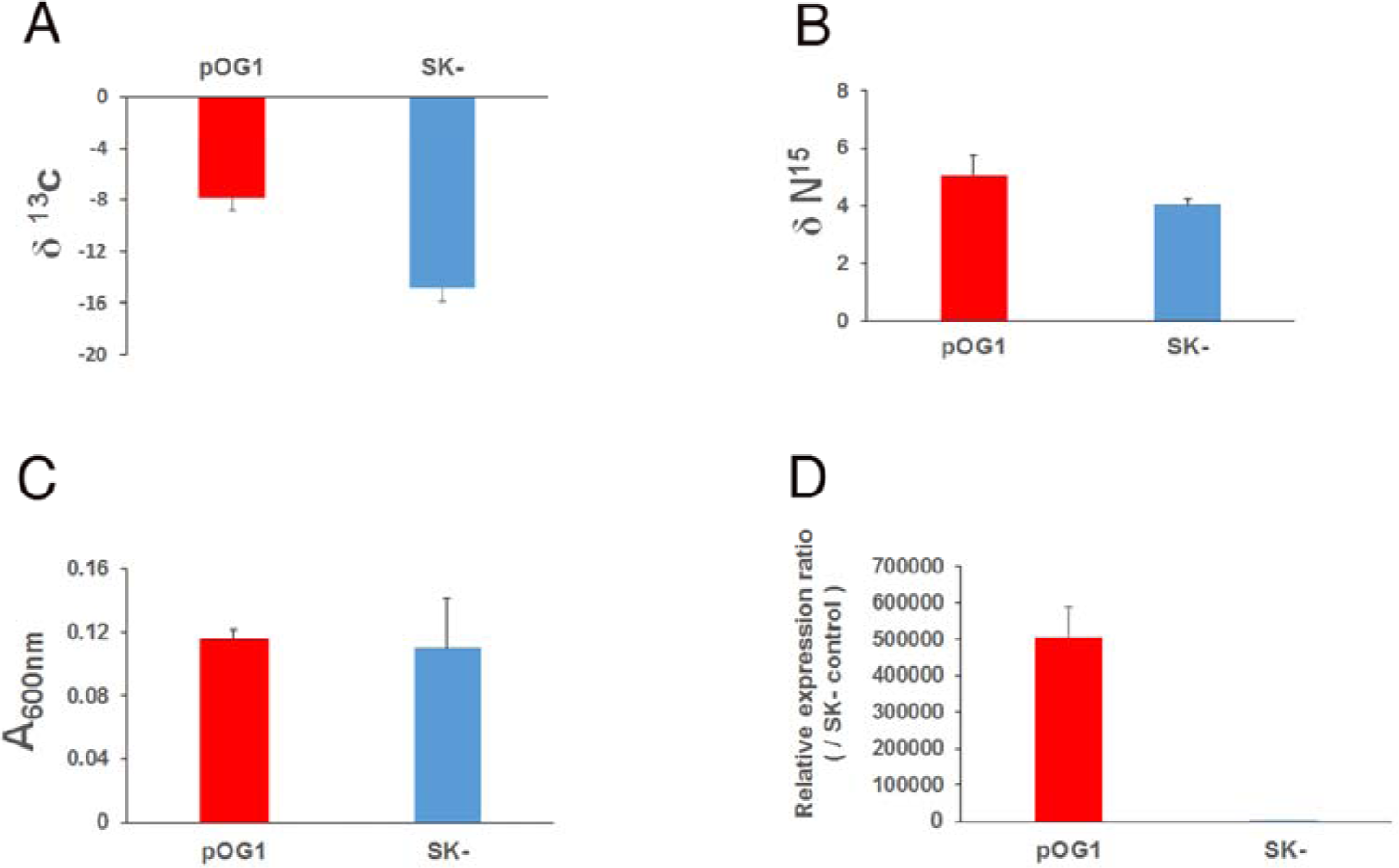
*OG1* gene expression in *E. coli* clones post-CO_2_ capture. **A.** δ_13_C. **B.** δN^15^. **C.** OD600nm post-CO_2_ capture. **D.** Relative gene expression of *OG1* in the ratio of gene expression in pOG1/pSK- clones in *E. coli* MG1655 cells. Statistical evaluations for δ_13_C are as follows (two tailed, and n=3): **A.** 0.015 (pOG1 vs pSK-; univariate General Linear Model). **D.** 0.008 (pOG1 vs pSK-, t test). The data were presented as average values with one standard deviation.

**Table 1.** Mutants from directed evolution.

| Name | GenBank<br>Accession no. | Site in the<br>OG1 gene | DNA Base mutation | Amino acid<br>substitution |
| --- | --- | --- | --- | --- |
| pOGDR1 | MF968946 | 322 | C>A | P108T |
| pOGDR2 | MF968952 | 237 | T>C | Nonsense mutation |
| pOGDR3 | MF968947 | 113 | A>G | N38S |
| pOGDR4 | MF968948 | 289 | A>G | I97V |
| pOGDR5 | MF968949 | 389,96 | A>C, T>C | H130P |
| pOGDR6 | MF968950 | 389 | A>C | H130P |
| pOGDR7* | JX462739 | 269 | A>G | Q90R |
| pOGDR10 | MF968951 | 169 | C>T | H57Y |

\*OGDR7 shares homology with *Homo sapiens isolate P75 mitochondrion*
(GenBank Accession no.: JX462739)

## Discussion

The subclone pOG1, identical to a known protein OK/SW-CL.16, shares homology with numerous proteins including Cytochrome c oxidase subunit III. It is a constituent of the multi-subunit cytochrome c oxidase, and part of the respiratory chain of mitochondria and aerobic bacteria^6,7^. It catalyzes the following reaction:

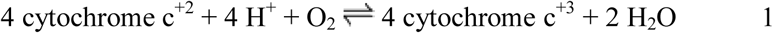

The utilization of the four protons above might be tied to the hydration of CO_2_ below, which leads to dissociation of protons and generation of bicarbonic acid.

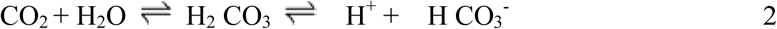

Bicarbonic acid could be assimilated as carbon source^8^, and accumulated protons could be utilized by OG1 protein in the aforementioned first reaction coupled with second reaction via proton traffic to produce water molecules without generating excessive osmotic pressure.

The pOG1 clone was initially isolated in nitrogen free media. Although acetylene reduction assays indicated no obvious nitrogen fixation activity with pOG1 clone, weak activity may arise endogenously in the host upon the potential regulation by the OK/SW-CL.16 protein that shares homology with PII uridylyl-transferase, which acts as regulator of the cellular nitrogen and carbon status in prokaryotes and plants^9^, such as NifA activity regulation and ammonium-dependent post-translational regulation of nitrogenase in *H. seropedicae*^10^.

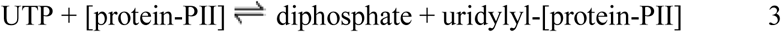

In response to nitrogen limitation, the uridylylated PII protein promotes deadenylylation of glutamine synthetase (GlnA), consequently activates the enzyme and stimulates NtrC-dependent promoters^11^.

The P108T substitution in pOGDR1 had profound impact on carbon fixing outcome in media with different pH values. The σ_22_ elements of the ^13^C chemical shift anisotropy for the deprotonated carboxyl groups indicate stronger hydrogen bonding of carbonyl oxygen in threonine than that in either proline or serine^12^. The electron-withdrawing inductive effect of the hydroxyl oxygen atom renders the hydroxyl oxygen in β-branched threonine less chemically active than its counterpart in serine, for instance, in the formation of hydrogen bonds. These properties will impact enzyme catalysis by the protein of OK/SW-CL.16 variant as a whole. The potentially non-casual connection between respiration chain or ATP’s effect and CO_2_ usage suggests that the mitochondrial genome may harbor other genes for CO_2_ capture. Another potential application of the *OG1* gene would be serving as a selective marker by growing on carbon deficient minimal media and replacing antibiotic selective markers in constructed plasmid.

Given the minimal nutrition it requires and potential regulation by soil ions and other substances on proton traffic, the pOG1 clone could be dispersed in barren lands to capture large amount of atmospheric CO_2_ to cope with escalating global warming crisis.

## Methods

### Cloning and Subcloning of the gene capable of CO2 capture

Chimeric DNA fragments encoding *S. cerevisiae* α-factor secretion signal and random peptides were generated from PCR amplifications of pPICZαA (Invitrogen, Carlsbad, CA, USA) linearized with BamH I. The two primers were: (1) GCGGATCCAAAAATGAGATTTCCTTCAA;(2)CGTCTAGATCAACCN(T/C)(G/ C/T)N(T/C)(G/C/T)ACAN(T/C)(G/C/T)N(T/C) (G/C/T)ACCAGATGGN(T/C)(G/C/T)N(T/C)(G/C/T) ACAN(T/C)(G/C/T)N(T/C)(G/C/T)AGATCTTTTCTCGAGAGATACCC. PCR reactions were performed using *Pfu* and *Taq* DNApolymerases (Takara, Dalian, China) in the presence of 2.5% pyrrolidone (Tokyo Kasei Kogyo Co. Ltd., Chuo-ku, Japan). After a preheating step at 94 °C for 2 min, PCR amplification was performed with 40 cycles of incubation at 95 °C for 20 s, 20 °C for 60 s and 72 °C for 30 s, and a final extension at 72 °C for 5 mins. PCR product was ethanol precipitated and dried. Double digestion with Xba I and BamH I was subsequently conducted for 5 h. pYES2/CT-ADH2 Vector^13^ was triple digested with *Bam*H I, *Xho* I and *Xba* I^14^, and then ligated with double digested PCR product at 6 °C overnight. Ligation mixture was electroporated to *E. coli* ORIGAMI cells (Novagen, Madison, WI, USA) at 12.5 kV/cm, followed by plating to nitrogen fixation media (M9 media with NH_4_Cl removed, and supplemented with 1% sucrose as carbon source)^15^. One colony appeared after growing at 30 °C for 3 days. The positive clone pOG was unable to grow after 3-month storage at the −20 °C freezers, and subsequently revived by 1-min UV irradiation in a laminar hood followed by plating to nitrogen fixation media described above. It was then successfully sequenced only after adding 5 mg/L Vitamin C to LB for cell propagation. pOG was subsequently amplified with Taq DNA polymerase with the following 2 primers: F0: GACGGTATCGATAAGCTTGATATCGAATTCATGACCCCTAACAGGGGC; PB2: ACTAGTGGATCCCCCGGGCTGCAGCTATGGTGAGCTCAGGTGATTGATACTCCTGATGC. After a preheating step at 94 °C for 3 mins, PCR amplification was performed with 35 cycles of incubation at 95 °C for 20 s, 42 °C for 45 s and 72 °C for 90 s, and a final extension at 72 °C for 5 mins. The PCR product was purified with the QIAquick gel extraction kit (QIAGEN, GmbH, Hilden, Germany) per manufacture’s instruction. Both PCR amplicons and SK- vector were double digested with *EcoR* I and *Bam*H I at 37 °C for 4 h, which were subsequently purified with the QiaQuick kit. Ligation was carried out at 16 °C overnight before electroporating into *E. coli* MG1655 cells. Positive clone pOG1 was identified after plasmid sequencing.

### _13_C-CO_2_ assimilation assays

pSK- clone and pOG1 clone in MG1655, and MG1655 cells were inoculated to LB respectively, and propagated at 37 °C at 120 rpm for 19 h. The cell pellets were washed once with sterile water and resuspended in sterile water. 4 OD (600 nm) of cells were inoculated to 50 ml CO_2_ capture media (0.02% YNB (Yeast Nitrogen Base, nitrogen free, Amresco, Inc., Solon, Ohio), 0.01 % ammonia sulfate, 0.04% yeast extract, pH 8.0 adjusted with NaOH) with ampicillin (60 mg/L) or no ampicillin for MG1655. Six layers of cheese cloth were used as caps to allow CO_2_ in. Vacuum desiccator was adapted for use with atmosphere inside. CO_2_ was released by hydrolysis of 6 mg urea-^13^C (99 atom %, Sigma Aldrich, Missori, USA) by 0.2 mg of urease from Jack Bean (Tokyo Chemical Industry Co. Ltd., Chuo-ku, Tokyo, Japan) in PBS (pH 7.4) in a total volume of 150 µl. The apparatus was sealed with Vaseline, and cells were cultured inside for 96 h at 30 °C. *E. coli* MG1655 cells was cultured outside of the device with ordinary airtight cap at the same temperature and with the same incubation time. Subsequently absorbance at 600 nm was measured. Bacteria were centrifuged for 10 min at 7084 x g, and the pellets were washed with 20-30 ml of sterile water and mixed well followed by centrifugation for 5 min at 7084 x g. Pellets were transferred to 1.5 ml Eppendorf tubes followed by drying at 60 °C for 1 h. For *S. cerevisiae*: Sample PYES2OG1 clone and control pYES2/CT/α-Factor clone in INVSc1 were divided into two groups with five replicates each. 5 OD (600 nm) of cells were inoculated to 50 ml CO_2_ capture media (0.02% YNB, 0.01 % ammonia sulfate, 0.04% yeast extract, pH 8.0 adjusted with NaOH, adding 1x amino acids concentrate of final concentration before use) with ampicillin (100 mg/L). 100 x amino acids concentrate (Ura^−^) contained 5 g Adenine,5 g leucine, 5 g tryptophan and 2.5 g histidine per liter. CO_2_ was released by hydrolysis of 6 mg urea-^13^C (99 atom %, Sigma Aldrich, Missori, USA) by 0.2 mg of urease from Jack Bean (Tokyo Chemical Industry Co. Ltd., Chuo-ku, Tokyo, Japan) in 10 mM PBS (pH 7.4) in a total volume of 150 µl. The rest is as described above.

### Determination of stable carbon and nitrogen isotope ratio

Experiments were carried out in the Environmental Stable Isotope Lab, Chinese Academy of Agricultural Sciences^16^. 2-4 mg of sample was weighed in a tin foil cup, and was subsequently introduced into the elemental analyzer (Vario PYRO cube, Elementar, Germany) through an automatic sampler, where the sample was burned and reduced into pure CO_2_ and N_2_. The gases were further diluted in a dilution apparatus, and measured in the stable isotope ratio mass spectrometer (IsoPrime100, Isoprime, England). The detailed running parameters were as follows: Elemental Analyzer: Burner temperature: 1020 °C; Reducing furnace temperature: 650 °C; He carrier gas flow rate: 230 mL/min. Dilution Apparatus: He dilution pressure: 4 bar; CO_2_ reference gas pressure: 8 psi; N_2_ reference gas pressure: 8 psi. Mass Spectrometer: CO_2_ reference gas was calibrated *via* USGS24 (δ^13^C_PDB_ = −16‰) through two point corrections, and the results were corrected by USGS24 and IAEA600 (δ^13^C_PDB_ = −27.5‰). N_2_ reference gas was calibrated *via* IAEA N1 (δ^15^N_air_ = 0.4‰), and the results were corrected by IAEA N1 and USGS43 (δ^14^N_air_ = 8.44‰).

### Liquid culture in carbon fixing media with or without yeast extract

Carbon fixing medium A contained 0.2 g/L YNB, 0.1 g/L ammonia sulfate, 0.4 g/L yeast extract, with sodium hydroxide adjusting pH equal to 8.0, supplemented with 100 μg/ml ampicillin. Medium B was as above but free of yeast extract. Two samples designated pOG1 and SK- in *E. coli* MG1655 were propagated overnight in LB medium with 100 μg/ml ampicillin at 37 °C and aerated by shaking at 220 rpm. Cultures were centrifuged for 10 min at 6000 rpm and washed with sterile ultrapure water twice after reaching logarithmic phase. The washed bacteria with OD_600nm_ of 0.2 were inoculated into carbon fixing medium with 100 μg/ml ampicillin. Each group had four replicates. OD_600nm_ was determined after incubation at 30 °C for 3 days (Supplementary Fig. S3).

### pH tolerance test

Carbon fixing medium contained 0.2 g/L YNB, 0.1 g/L ammonia sulfate, 0.4 g/L yeast extract, with sodium hydroxide adjusting pH equal to 5.0, 6.0, 7.0, 8.0 and 9.0 respectively. Bacterial cultures were propagated overnight in LB medium with 100 μg/ml ampicillin at 37 °C with shaking at 180 rpm. Cultures were centrifuged for 5 min at 6000 rpm and washed with sterile ultrapure water twice after reaching logarithmic phase. The washed bacteria with OD_600nm_ of 0.01 were inoculated into carbon fixing media with 100 μg/ml ampicillin, Each sample had three replicates with 5 ml liquid media in the tubes. OD_600nm_ was determined after incubation at 30 °C for 12 h (Supplementary Fig. S4).

### Construction of a *S. cerevisiae/E. coli* Shuttle Vector

pOG1 in *E. coli* was propagated overnight in LB medium with 100 μg/ml ampicillin at 37 °C at 220 rpm. *OG1* gene was amplified with AlphaOG1 primer 5′-TCTCTCGAGAAAAGAATGACCCCTAACAGGG and 3OG1 primer 5′-GATCTAGACTATGGTGAGCTCAGG. The PCR mixture (50 μl) consisted of 5 µl of 10x PCR Buffer (Mg^2+^ plus), 1 µl of template, 1.5 µl of 10 mM dNTP mix, 1µl of 10 mM upstream primer, 1 µl of 10 mM downstream primer, 0.3 µl of Taq enzyme (5 U/µl), and sterilized distilled water up to 50 μL. PCR conditions were as follows: 2 min initial denaturation at 94 °C; then 35 cycles of 95 °C for 20 s, 55 °C for 45 s, 72 °C for 90 s; and finally 60 s at 72 °C. PCR products were purified by Gel&PCR purification kit (Promega, Madison, WI, USA). The PCR amplicons and pYES2/CT/α-factor (Changsha Yingrun Biotechnology Co., China) were digested with *Xho* Ↄ and *Xba* Ↄ for 4 hours at 37 °C, followed by inactivation at 70 °C for 10 minutes and ligation at 16 °C. The ligation products were purified via Gel&PCR purification kit, and electroporated into *E. coli* DH5α, which was then plated on a LB agar plate supplemented with 100 μg/ml ampicillin and incubated overnight at 37 °C. The positive colonies were verified by PCR followed by sequencing. The resultant plasmid was designated pYES2OG1.

### Directed evolution

pOG1 clone in *E. coli* was propagated overnight in LB medium with 100 μg/ml ampicillin at 37 °C at 220 rpm. *OG1* gene was amplified with OG1F0 primers 5′-TCGAATTCATGACCCCTAACAGGG and OG1R0 primers 3′-AGTGGATCCCCCGGGCTGCAGCTATGGTGAGCTCAGG. Instant Error-prone PCR Kit was a product of Beijing Tianenze Gene Technology Co., Ltd. (Beijing, China)^17^. The Error-Prone PCR mixture consisted of 5 µl of 10x Error-Prone PCR mix, 5 µl of 10x Error-Prone PCR proprietary dNTP, 5 µl of 5mM MnCl_2_, 1 µl of template, 0.5 µl of 10µM upstream primer and downstream primer each, 0.6 µl of Taq DNA polymerase (5 U/µl), and sterilized distilled water up to 50 μl. Error-Prone PCR conditions were as follows: 40 cycles of 98 ° for 10 s, 55 °C for 55 s, 72 °C for 60 s. PCR products were purified by Gel&PCR purification kit. Both PCR amplicons and pSK- vector were double digested with *Eco*R I and *Bam*H I at 37 °C for 4 h, followed by inactivation at 80 °C for 15 minutes and ligation at 16 °C. The ligation products were purified and recovered by Gel&PCR purification kit, and electroporated into *E. coli* DH5α cells, followed by plating on a LB agar plate supplemented with 100 μg/mL ampicillin and incubating overnight at 37 °C. The positive colonies were verified by colony PCR followed by sequencing. The resultant plasmid was named pOGDR1, pOGDR2, pOGDR3, pOGDR4, pOGDR5, pOGDR6, pOGDR7, pOGDR10. After an initial one-sample screening via isotopic experiment, clones with higher carbon fixing activities were further investigated in subsequent experiments.

### Real-time fluorescent quantitative PCR

The experiment was contracted with FitGene BioTechnology Co., LTD (Guangzhou, China)^18^. Briefly, an aliquot of the six samples were collected at the end of a 4-day carbon fixing experiment. RNA was extracted using RNAprep Pure Cell/Bacteria Kit according to the instructions of the manufacturer (TIANGEN Biotech Co., Ltd, Beijing, China). The RNA purity was determined by UV spectrophotometer SMA4000 (Merinton, China). The RNA was added to the gDNA adsorption column and centrifuged at 10000 g for 1 min at room temperature for removal of genomic DNA. The RNA was thermal denatured at 65 °C, and immediately cooled on ice for 2 min, and used as template for reverse transcription. Synthesis of cDNA was performed in 10 μL reactions per manufacturer’s instructions. After the reaction, the cDNA was diluted 5 times with sterilized deionized water and kept at −20 °C. Real-time fluorescent quantitative PCR was then conducted. 20 μl PCR reactions containing template cDNA, 1x Bestar^®^ SybrGreen qPCR Mastermix, 1x ROX, 0.2 μM of each primer, and sterilized distilled water were performed in an ABI7500 thermal cycler (Life Technologies, Carlsbad, CA, USA). The cycling conditions comprised 2 min polymerase activation at 95 °C, 45 cycles at 95 °C for 10 seconds, 60 °C for 34 s (Fluorescence signal acquisition) and 72 °C for 30 s. In the end of the cycle, melting curve from 60 °C to 98 °C was obtained. The specificity of the amplifications was verified by melt curve analysis. Gene expression levels were normalized (ΔΔCt analysis) to 16S rRNA gene expression levels from the same sample. Gene expression from relative real-time quantitative PCR experiment was determined using 2^−ΔΔCT^ method. All data are expressed as the means ± standard deviation.

### Treatments with carbonic anhydrase inhibitor acetazolamide on CO2 capture

pSK- clone and pOG1 clone in *E. coli* MG1655 were each divided into two groups according to the presence (+) or absence (−) of 50 µg/ml acetazolamide (Sigma, Missori, USA) in the media. Each group had three duplicates. CO_2_ capture experiment was conducted as aforementioned.

### Statistical analyses

Statistical analyses were performed using SPSS 22.0. The experiments were evaluated using the Univariate General Linear Model or independent samples-T test when data were normal distributed or approximately normal distributed after examining with Shapiro-Wilk tests. The alpha level for all tests was 0.05. Games-Howell post hoc tests were conducted when equal variance was not assumed.

The OG sequence in the original positive clone has been deposited in GenBank with the accession number of KX255659. *OG1* sequence is identical to a GenBank sequence with the accession number NC_012920, ranging from 9251 to 9655 bp. Its encoded protein OK/SW-CL.16 was previously deposited in GenBank with the accession number of BAB93516.

### Data Availability

All data generated or analyzed during this study are included in this published article and its Supplementary Information file.

## Acknowledgements

This work was supported by grants from the Guangzhou Science and Technology Program (201804010328); Guangdong Science and Technology Program (2016B020204001, 2008B020100001); Guangdong Natural Science Foundation (S2011010004264); The National Natural Science Foundation of China (30370799) and The Science and Technology Transformation Program of Sun Yat-sen University (2019) to Q. Liu; Open Fund of Laboratory (20160215) at Sun Yat-sen University of China to X. Zhu; The National Natural Science Foundation of China (31330080), China Agriculture Research System (47) to J.H.; The National Natural Science Foundation of China (21601209) to Y.C.W. We thank technical help and suggestions from M. Jia, J. Zhou, Q. Zhi, Z. Tan, G. Yu, S. Li and S. Wang. Manuscript proofreading by Y. Shi is appreciated.

## References

1. Yu, K. M. K., Curcic, I., Gabriel, J. & Tsang, S. C. E. Recent Advances in CO2 Capture and Utilization. ChemSusChem 1, 893–899 (2008).

2. Macdowell, N. et al. An overview of CO2 capture technologies. Energ. Environ. Sci. 3, 1645–1669 (2010).

3. Togashi, M. et al. Interaction of alpha-actinin-4 with class I PxxP motif-containing OK/SW-CL.16 protein. Nephron Exp. Nephrol. 107, e65–72 (2007).

4. Chey, W. D., Wong, B. C. & Practice Parameters Committee of the American College of Gastroenterology. American College of Gastroenterology guideline on the management of Helicobacter pylori infection. Am. J. Gastroenterol. 102, 1808–1825 (2007).

5. Malfertheiner, P. et al. Current concepts in the management of Helicobacter pylori infection: the Maastricht III Consensus Report. Gut 56, 772–781 (2007).

6. Belevich, I., Verkhovsky, M. I. and Wikström, M. Proton-coupled electron transfer drives the proton pump of cytochrome c oxidase. Nature 440, 829–832 (2006).

7. Michel, H. Cytochrome c oxidase: catalytic cycle and mechanisms of proton pumping--a discussion. Biochemistry 38, 15129–15140 (1999).

8. Cramer, M. D., Lewis, O. A. M. & Lips, S. H. Inorganic carbon fixation and metabolism in maize roots as affected by nitrate and ammonium nutrition. Physiol. Plantarum 89, 632–639 (1993).

9. Garcia, E. & Rhee, S. G. Cascade control of Escherichia coli glutamine synthetase. Purification and properties of PII uridylyltransferase and uridylyl-removing enzyme. J. Biol. Chem. 258, 2246–2253 (1983).

10. Noindorf, L. et al. Role of PII proteins in nitrogen fixation control of Herbaspirillum seropedicae strain SmR1. BMC Microbiol. 11, 8 (2011).

11. Perlova, O., Nawroth, R., Zellermann, E. M. & Meletzus, D. Isolation and characterization of the glnD gene of Gluconacetobacter diazotrophicus, encoding a putative uridylyltransferase / uridylyl-removing enzyme. Gene 297, 159–168 (2002).

12. Gu, Z., Zambrano, R. & Mcdermott, A. Hydrogen bonding of carboxyl groups in solid-state amino acids and peptides: comparison of carbon chemical shielding, infrared frequencies, and structures. J. Am. Chem. Soc. 116, 6368–6372 (1994).

13. Wang, S. et al. Isolation of oligopeptides targeting Ras-Raf interactions via reverse two hybrid assays. Biotechnology 24, 58–63 (2014).

14. Xu, Z. et al. Isolation of novel sequences targeting highly variable viral protein hemagglutinin. MethodsX 2, 64–71 (2015).

15. Sambrook, J., Maniatis, T. & Fritsch, E. F. Molecular cloning: A Laboratory Manual (Cold Spring Harbor Laboratory Press, 1989).

16. Johnson, C. A., Stricker, C. A., Gulbransen, C. A. & Emmons, M. P. 2018, Determination of δ^13^C, δ^15^N, or δ^34^S by isotope-ratio-monitoring mass spectrometry using an elemental analyzer: U.S. Geological Survey Techniques and Methods, book 5, chap. D4, 19 p.

17. Leung, D. W., Chen, E. & Goeddel, D. V. A method for random mutagenesis of a defined DNA segment using a modified polymerase chain reaction. Technique 1, 11–15 (1989).

18. Jothikumar, N. & Griffiths, M. W. Rapid detection of Escherichia coli O157:H7 with multiplex real-time PCR assays. Appl. Environ. Microbiol. 68, 3169–3171 (2002).

